# Robust detrending, rereferencing, outlier detection, and inpainting for multichannel data

**DOI:** 10.1101/232892

**Authors:** Alain de Cheveigné, Dorothée Arzounian

## Abstract

Electroencephalography (EEG), magnetoencephalography (MEG) and related techniques are prone to glitches, slow drift, steps, etc., that contaminate the data and interfere with the analysis and interpretation. These artifacts are usually addressed in a preprocessing phase that attempts to remove them or minimize their impact. This paper offers a set of useful techniques for this purpose: robust detrending, robust rereferencing, outlier detection, data interpolation (inpainting), step removal, and filter ringing artifact removal. These techniques provide a less wasteful alternative to discarding corrupted trials or channels, and they are relatively immune to artifacts that disrupt alternative approaches such as filtering. *Robust detrending* allows slow drifts and common mode signals to be factored out while avoiding the deleterious effects of glitches. *Robust rereferencing* reduces the impact of artifacts on the reference. *Inpainting* allows corrupt data to be interpolated from intact parts based on the correlation structure estimated over the intact parts. *Outlier detection* allows the corrupt parts to be identified. *Step removal* fixes the high-amplitude flux jump artifacts that are common with some MEG systems. *Ringing removal* allows the ringing response of the antialiasing filter to glitches (steps, pulses) to be suppressed. The performance of the methods is illustrated and evaluated using synthetic data and data from real EEG and MEG systems. These methods, which are are mainly automatic and require little tuning, can greatly improve the quality of the data.

## 1 Introduction

The very weak brain signals picked up by electroencephalography (EEG) or magnetoencephalography (MEG) have to compete with multiple sources of noise and artifact within the body, the environment, and the sensors or electrodes. Of particular concern are electrode-specific or sensor-specific sources, because they cannot be suppressed by combining channels linearly as in ICA, beamforming or other linear techniques (Parra et al., 2005; Debener et al., 2010). In EEG they include *slow drifts* at the electrode/gel/skin
interface (Huigen et al., 2002; Kappenman and Luck, 2010), and in MEG the large amplitude steps that result from a slip in the flux-lock loop (Gross et al., 2013), as well as various other glitches of diverse nature. We will use the terms “artifact” and “noise” interchangeably as the distinction between them is not well defined.

Many techniques have been proposed to eliminate or palliate artifacts, some of them well-established and included in standard guidelines and processing pipelines These include temporal and spatial filtering, detrending, regression, rereferencing, rejection of corrupt data, and spatial interpolation. However, many of these methods are prone to failure for certain combinations of artifact, and in some cases they may make things worse. As an example, *high-pass filtering*, a standard method to deal with drifts, is highly sensitive to the presence of temporally-localized glitches that trigger ringing of the filter. Here we revisit the issue of preprocessing, and propose a methodology of robust estimation of regression parameters, including methods for *detrending, outlier detection, inpainting, rereferencing, step removal*, and *ringing artifact removal*. These methods have in common that they deal with artifacts that are not easily removed with standard spatial filtering techniques.

We first review some important artifacts that affect EEG and MEG data.

### Drifts in EEG

The EEG signal often rides upon a slow drift signal produced at the skin / electrolyte / electrode interface (Huigen et al., 2002). Figure 1 shows the time course of a typical sample of EEG data after removal of the mean from each channel. Each channel seems to follow its own drift pattern, albeit with some apparent inter-channel correlation. Slow drift can be minimized at the source, by piercing or abrading the outermost layer of the skin, the stratum corneum (Stjerna et al., 2010; Vanhatalo et al., 2005), but not all experimenters or subjects welcome this solution. Slow drifts may mask genuine cortical activity in the very low frequency range (Vanhatalo et al., 2005), and furthermore, if the data are epoched, the drift may misleadingly appear as a pattern reproducible over trials, a tendency that may be further reinforced by component analysis techniques that emphasize repeatable components (de Cheveigné and Parra, 2014). Drifts are usually dealt with by high-pass filtering or detrending, but unfortunately those methods themselves may create new artifacts as discussed in the next paragraphs.

**Figure 1:**
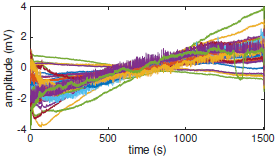
Sample of 40-channel EEG data with slow drifts. Data were acquired with a BioSemi system at a rate of 2048 Hz in the calibration phase of an experiment investigating auditory perception and brain state. The mean of each channel was subtracted before plotting.

### High-pass filtering artifacts

High-pass filtering is effective to attenuate slow drifts (typical cutoffs range from 0.1 to 2Hz) but it raises several issues. Most obvious is the removal of slow cortical activity, whether spontaneous (Vanhatalo et al., 2005), or stimulus-evoked (Southwell et al., 2017). Processing that blinds the investigator to such cortical activity is not ideal. Another problem, less often appreciated, is that transient features too may be distorted due to the filtering (Acunzo et al., 2012; Vanhatalo et al., 2005). Figure 2 (top left) shows how a unipolar pulse (0.5 s duration) is affected by typical high-pass filters (Butterworth design) of various orders and cutoff. The pulse is attenuated relative to the original signal, and is followed by multipolar excursions that are *purely artifactual*, with a morphology that depends on the cutoff frequency, order, and type of filter. A transient cortical event would incur similar distortion, which is worrisome as the additional excursions might masquerade as distinct neural events. The point has been repeatedly made (Acunzo et al., 2012; Tanner et al., 2015; Widmann et al., 2015; Tanner et al., 2016), but it escapes consideration in many studies.

In the same vein, Figure 2 (top right) shows how a step (0.5 s rise time), such as might occur in the cortical response to a stimulus onset (Southwell et al., 2017), is affected by the same filters. The sustained plateau is lost and replaced by spurious signal excursions of both signs that would completely obfuscate the pattern of cortical electrical response. An offset would trigger a similarly misleading “offset response” (not shown). Zero-phase filtering is sometimes recommended as conducive to less distortion, but it introduces an additional issue. Figure 2 (top right, bottommost lines) shows the response of zero-phase filters to the same step. The presence of spurious deflections is now made worse by the fact that some of them occur *before* the event that triggered them. Applied to neural data, non-causal filters may confuse our understanding of the sequence of events within the brain and their causal relations.

Figure 2 (bottom) illustrates a high-pass filter response to a descending ramp (analogous to the EEG data in Fig. 1). After a transient response (due to the implicit step at the onset) the signal is close to zero, as desired. However the transient fluctuations take time to subside, especially for a filter with a lower cutoff and/or higher order. They are of particular concern if the high-pass filter is applied *after* cutting the data into epochs (not recommended), as the fluctuations can easily be mistaken for a reproducible neural response. Such fluctuations may be reduced by including more data at the beginning of each epoch, to give time for the transient to subside, however the response of an infinite impulse response (IIR) filter never completely dies away. Together, these issues provide strong motivation to consider alternatives to high-pass filtering.

**Figure 2:**
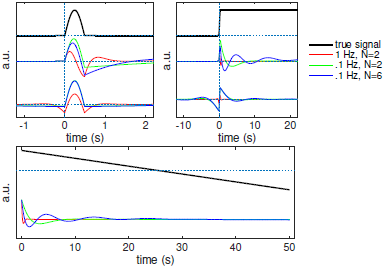
Top: pulse (left, black) and step (right black) signals and corresponding filter outputs for 3 different high-pass filters. Lowermost curves are for non-causal filters (Matlab `filtfilt` function). Bottom: descending ramp (black) and filter output for 3 different Butterworth high-pass filters with cutoff frequency and order (N) indicated in the legend.

### Detrending artifacts

In *detrending*, a smooth function, for example a low order polynomial, is fit to the data and then subtracted from it. Figure 3 (top left) shows one channel of an EEG signal contaminated with a slow drift, together with a polynomial fit of order 10 (red), and Figure 3 (top right) shows the same signal after subtraction of the fit. Slow fluctuations have disappeared, leaving rapid fluctuations on several time scales. The order of the polynomial controls the scale of the fluctuations that are removed.

Detrending is unfortunately sensitive to the presence of *glitches*, that are common in EEG and MEG. Figure 3 (middle left) shows the same data to which a glitch has artificially been added. The fit (red) is affected by the glitch, with the result that the detrended signal now shows high-amplitude fluctuations over an interval that extends outside the glitch-contaminated part (Figure 3 middle right). However, if the extent of the glitch is known, the fit can be restricted to the intact part (indicated by the grey bar). Figure 3 (bottom left, red) illustrates such a fit, and Figure 3 (bottom right) shows resulting detrended data. The glitch is of course still present, but the rest of the data are nicely detrended without spurious fluctuations. This approach is described in more detail in this paper.

**Figure 3:**
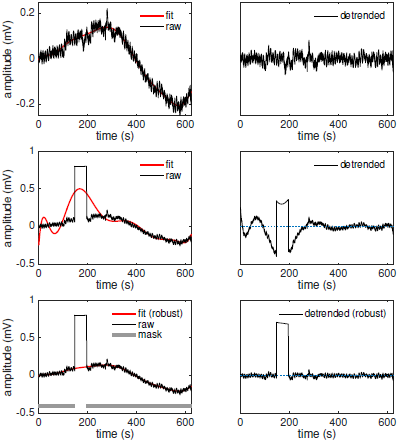
Top left: sample of EEG signal (black) and order-10 polynomial fit (red). Right: detrended data. Middle left: same EEG data as top with an artificial glitch (black) and polynomial fit (red). Right: “detrended” data. Bottom: same data with robust polynomial fit (red). The fit was weighted using the weighting function (mask) symbolized in grey. Right: detrended data. Data are from the same dataset as Figure 1.

### Rereferencing

EEG measures the potential at each electrode *relative* to some other electrode (Picton et al., 2000). The signal recorded on any channel obviously depends on the position of the reference electrode on the skull, and any noise introduced at the reference electrode (e.g. contact noise) contaminates all measured signals. To reduce the dependency on reference electrode position it is common to rereference to the average, i.e. subtract from each electrode the average over all electrodes (Nunes and Srinivasan, 2006; Keil et al., 2014) or some other linear combination (Yao, 2001). Noise at the physical reference electrode is cancelled, but noise on any other electrode is now injected, via the average, into *all* channels. This paper suggests a means to mitigate this issue.

### Glitches

EEG and MEG are susceptible to temporally-localized glitches, due to muscle artifacts or motion of the electrodes or leads. Channel-specific artifacts (that affect one rather than multiple electrodes) are particularly troublesome as they cannot be suppressed by linear techniques without effectively discarding the contaminated channel (de Cheveigné, 2016). They can interfere with data-driven linear techniques, as illustrated in Figure 3 (middle), and it is useful to identify them prior to other analysis.

Channel specific artifacts are usually uncorrelated across channels, in contrast to genuine brain activity that tends to be correlated across channels because of current spread (de Cheveigné and Simon, 2008). This suggests a way to *detect* them: (a) project a channel on the subspace spanned by all other channels, (b) estimate the statistics of the residual (channel minus projection), (c) find the outliers of this distribution. However this scheme runs into a practical difficulty: artifacts on other channels may corrupt the projection, and thus wrongly trigger the detection of artifacts on the channel of interest. To be effective, the algorithm should track the multiple combinations of intact/corrupt channels that can occur in real data, so as to always project on intact data. This paper describes such an algorithm.

*Inpainting* is a term used in the field of image processing to designate the process by which a missing portion of an image is replaced using a model constrained by the intact parts (Bertalmio et al., 2000), and the same idea has been used for other signals such as audio (Adler et al., 2012). Here we apply the concept to electrophysiological data. supposing the extent of an outlier or glitch (as in Figure 3 middle) is known (for example from the previous algorithm), it can be replaced using a model constrained by intact data. This paper proposes such a process. Inpainting and outlier detection are closely related.

### MEG steps

MEG has the advantage over EEG that it is not prone to the slow drift that arises at the electrode-skin contact in EEG. Slow nuisance signals may be present, due to environmental sources such as moving vehicles or machinery, but they are usually correlated across sensors and can be projected out of the data with linear methods. However, MEG is prone to sudden changes in the operating point of the flux lock loop associated with the SQUID sensors (Oswal et al., 2016; Gross et al., 2013), that produce large amplitude steps, against which high-pass filtering is particularly ineffective. This paper proposes a reliable method to remove these steps.

### Antialiasing filter artifacts

An event such as a flux jump (as mentioned above) or stimulus artifact (for example as occurs in cochlear or deep brain stimulation) necessarily triggers a ringing artifact due to the antialiasing filter that precedes conversion of the analog EEG or MEG signal to a digital representation. The ringing extends beyond the interval containing the event, and may remain after the event has been removed (as in the case of MEG steps). This paper proposes a method to address this issue.

In summary, electrophysiological data are plagued by many artifacts that interfere with the interpretation of the data, and that may reduce the effectiveness of processing algorithms needed to assist with this interpretation. Classic methods such as filtering are particularly prone to artifacts and signal distortion effects. This paper describes a panoply of alternative techniques to deal with these issues. These techniques have in common that they target artifacts that are not easily removed with standard spatial filtering techniques such as ICA. They are complementary with those techniques: once artifacts have been suppressed, techniques such as ICA may be more effective.

## 2 Methods

### 2.1 Signal models

We use three signal models. The first and last are applicable to a single channel independently from others, the second is applicable to multiple channels taken together.

For the first signal model, the data of channel *n* of a data matrix are expressed as:

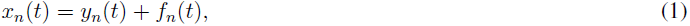

where *f_n_*(*t*) is the projection of *x_n_*(*t*) on some basis of functions (for example low-order polynomials) chosen to represent trends, and the residual *y_n_*(*t*) is a “detrended” signal. We assume that this model holds for all values of t except those for which a weighting function *w_n_*(*t*) is zero.

The second model is best described by first defining a simpler model that states that each channel can be expressed as a weighted sum of the other channels:

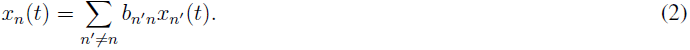

Equation 2 implies linear dependency between channels (rank of the data <*N*) but is a stronger condition. The idea expressed by this model is that no single channel is “indispensable”, as any channel can be reconstructed from the other channels. We will further assume that there are multiple ways to reconstruct the channel, based on different subsets of the other channels, i.e. eq. 2 also holds for subsets of {*n′* ≠ *n*}.

This simple model is now modified by assuming the presence of temporally sparse glitches, labeled by zero values of a weighting matrix *w_n_*(*t*). This may invalidate the baseline model (eq. 2), however, we make three assumptions. The first is that for every t such that channel *n* is valid (i.e. *w_n_*(*t*) ≠ 0), there exists some subset *O_n_* of valid channels {*n*′ ≠ *n*} for which eq. 2 holds. In other words, glitch-free parts of channel *n can* be reconstructed based on the other channels, although the “recipe” to do so might be different at different *t*, i.e. there are several such subsets 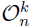. The one that is valid at *t* is notated as 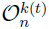. The second assumption is that at least one of these subsets 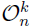 is also valid *during*the glitch that affects channel *n*. In other words it is possible to fix samples within the glitch-contaminated segment based on intact channels. The third assumption is that there are enough intact data so that all the required coefficients {*b_nn′_*}*_k_* in eq. 2 can be estimated reliably. These assumptions are plausible if the glitches are temporally sparse and don’t occur at the same time on all channels.

For the third model, each channel is expressed as:

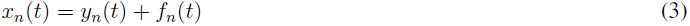

where *f_n_*(*t*) is now *piecewise constant*. This model will allow us to address steps in MEG data.

### 2.2 Robust detrending

This algorithm uses the first model. The slow trend on each channel *n* is estimated robustly using a weighting matrix *w_n_*(*t*), and then subtracted. The weighting matrix can be predetermined or else estimated iteratively with the following algorithm:

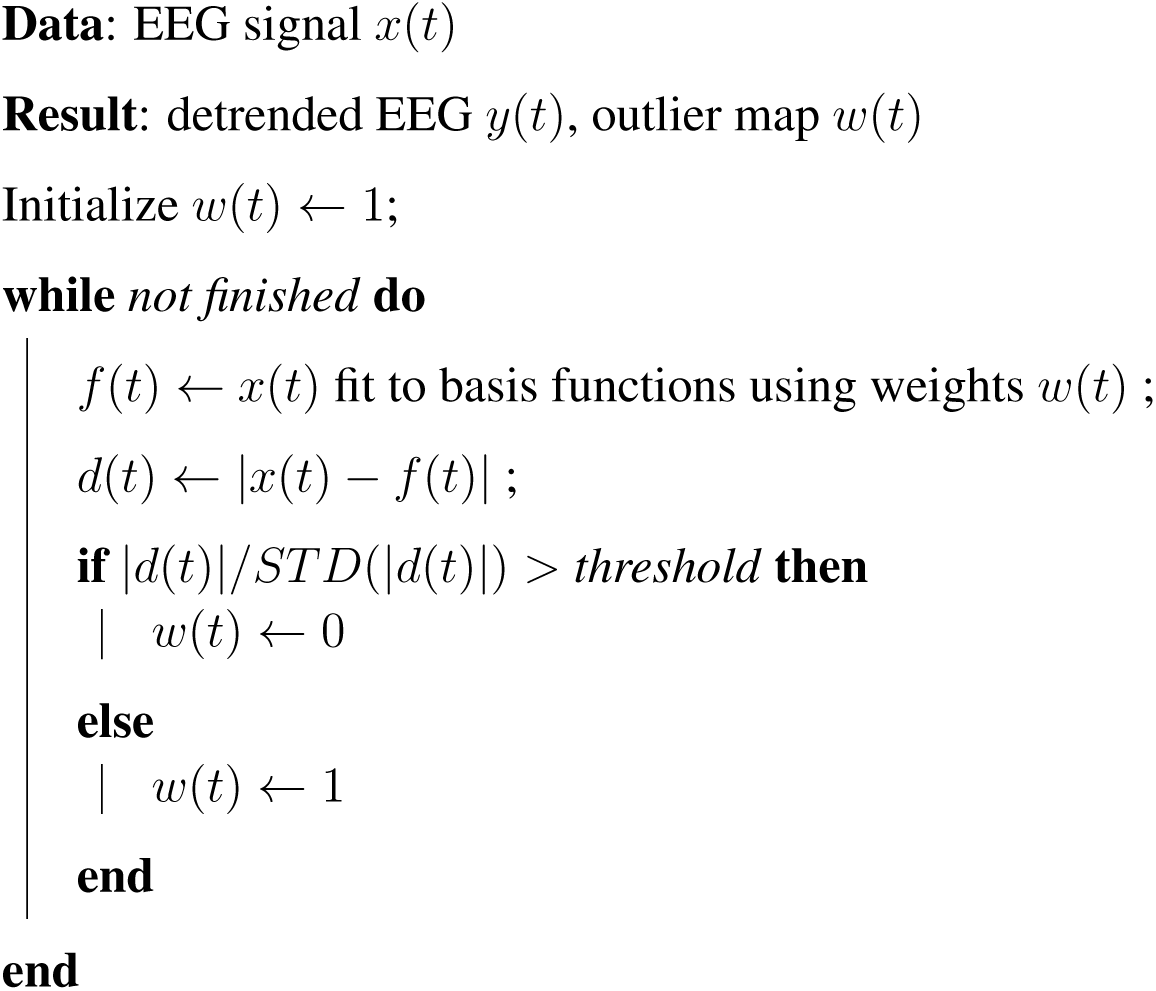

At each iteration the data are projected onto the basis functions, statistics of the residual *d*(*t*) are estimated, and samples larger than a threshold are flagged as outliers to be discounted in the next iteration. The loop is terminated after a few iterations, or if *w*(*t*) does not change between iterations. The weights *w*(*t*) can be initialized to values other than 1 if prior knowledge is available. A quality-of-fit score can be defined as 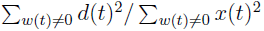. For multichannel data the algorithm is applied independently to each channel.

### 2.3 Inpainting

This algorithm uses the second signal model to estimate missing portions of multichannel data *x_n_(t)* (labeled by zero values of a weighting matrix *w_n_*(*t*)). According to that model, the signal *x_n_*(*t*) of channel *n* can be approximated for every *t* as a weighted sum 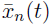) of a subset 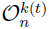 of the other channels. The subset may differ for different *t* according to which other channels are intact at that time. The algorithm estimates projection parameters based on intact portions of the data, and applies them to reconstruct the corrupted portions. More precisely:

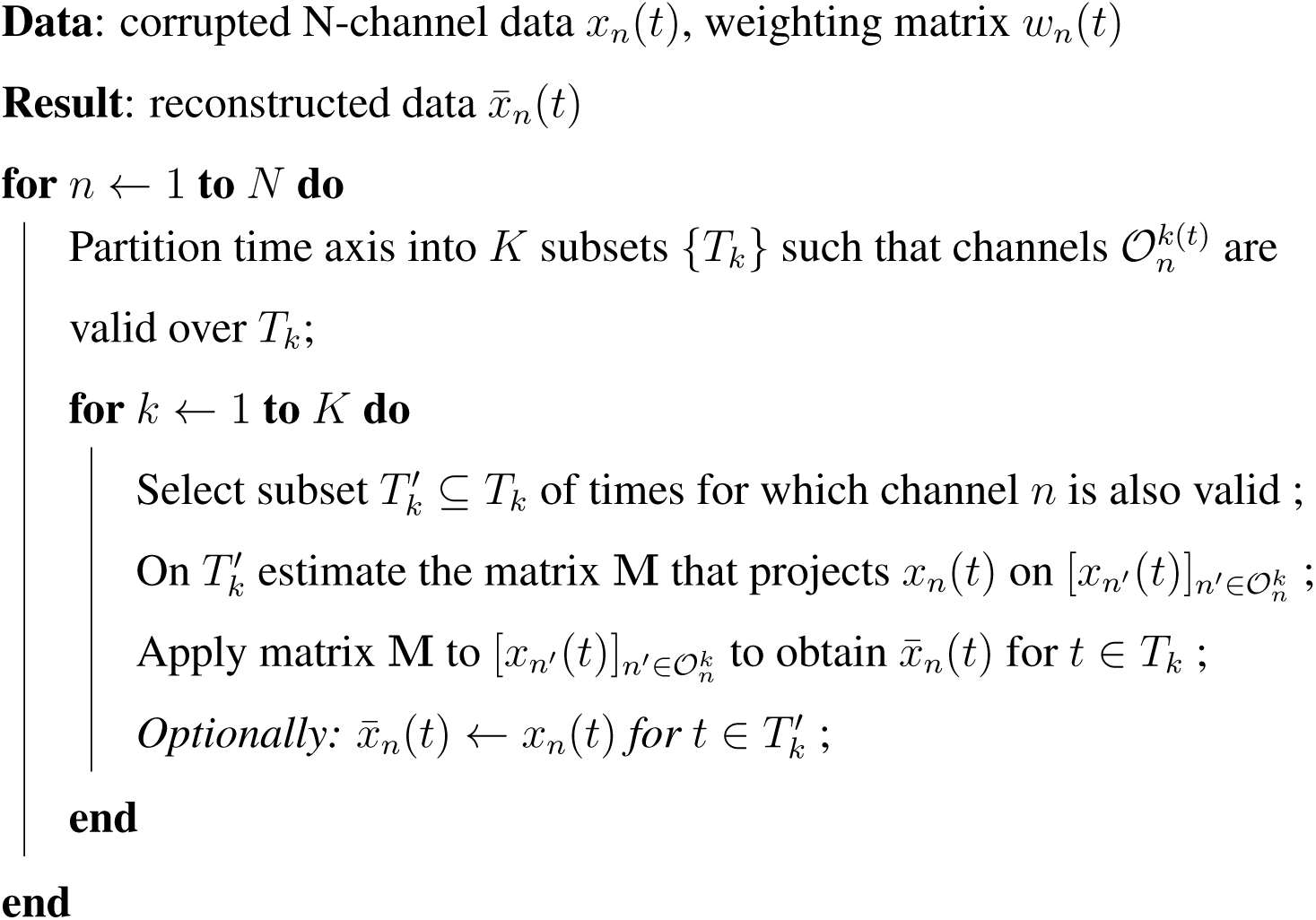

In brief, corrupt values of each channel are reconstructed from channels that are intact at that time. subsets *T_k_* or 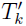 do not need to be formed of contiguous samples, but the subset 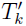 must be large enough to reliably estimate the projection matrix M. If the step labeled as *optional* is included, the output 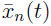 is identical to the original signal except during the glitch. If that step is omitted, the output is everywhere equal to a weighted sum of other channels (*n′* ≠ *n*), i.e. all output samples may differ from the input. The rank of the data is not usually reduced by processing.

### 2.4 Outlier detection

The previous algorithm requires a map *w_n_*(*t*) indicating which parts of the data are corrupt. If that information is missing it can be derived from the data using an adapted version of the algorithm. Starting with an initial estimate of the weights *w_n_*(*t*), for example all ones, each channel n is approximated on the basis of other channels *(n′ ≠ n)*. The approximation 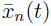 is then compared to the actual signal *x_n_*(*t*), and the times at which the mismatch exceeds a threshold are flagged as corrupt. More precisely:

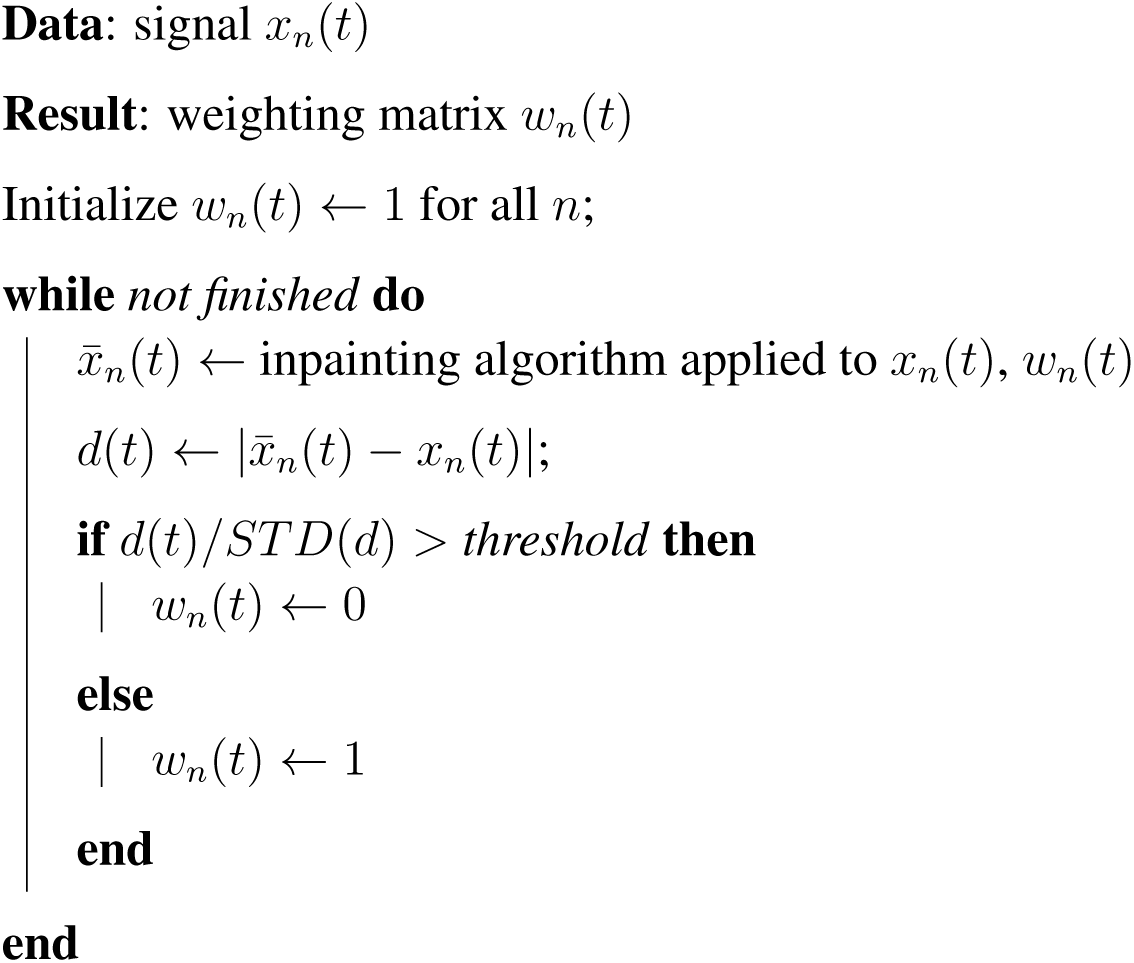

The algorithm is iterated a few times to refine the outlier map *w_n_(t)*. The value of *threshold* is not critical, a value of 1 is usually adequate. A quality-of-fit score can be defined as 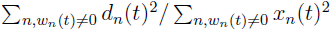.

### 2.5 Robust rereferencing

It is common in EEG to rereference the data by subtracting the average of signals over all channels *m(t) = (1/N) ∑_n_ x_n_*(*t*). As explained in the Introduction, a glitch on one channel can corrupt this average, and thus contaminate all channels. This can be avoided by replacing *x_n_*(*t*) in the formula for the mean by 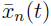 obtained from the inpainting algorithm. Alternatively, a simple expedient is to replace the mean over channels by a weighted mean: *m*(*t*) = ∑*_n_ w_n_*(*t*)/*x_n_*(*t*)/ ∑*_n_ w_n_*(*t*). Analogous schemes for robust rereferencing were proposed by Lepage et al. (2014); Bigdely-Shamlo et al. (2015).

### 2.6 Step removal

This algorithm uses the third signal model (piecewise constant) to locate and remove large amplitude steps such as SQUID jumps commonly observed in MEG data (Gross et al., 2013). The position *t*_0_ of a single jump in a signal *x(t)* of duration T can be found reliably as:

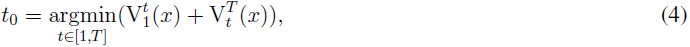

where 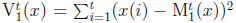 is the sum of squared deviations from the mean 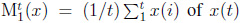 of *x*(*t*) over [1,*t*]. As described, the algorithm suffers from a slight bias towards choosing a step position close to either extremity (0 or *T*). This is because the variance of V increases as the number of samples used to calculate it decreases, making a spurious minimum in eq. 4 more likely. To counteract this trend and avoid trivial splits, it is convenient to restrict the search to the interval [*T_g_, T* − *T_g_*] where *T_g_* is a guard value. Likewise it is convenient to set a threshold on the ratio 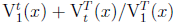 below which small amplitude “jumps” are ignored. Multiple jumps are located by recursing on each side of *t*_0_. The jumps are then removed from the data by subtracting the mean of the data between jumps. *Robust* step removal can be implemented by including a weighting term *w*(*t*) in eq. 4.

### 2.7 Ringing removal

After a step is removed by the previous algorithm, there remains a ringing artifact due to the step response of the antialiasing filter. If the characteristics of that filter are known, the ringing artifact can be removed by feeding a step to the same filter, and subtracting the response. A simple expedient to find the appropriate filter is to apply the Steiglitz-McBride iteration method (Steiglitz and Mcbride, 1965) (Matlab function `stmcb`) to a short segment of data following the step, yielding numerator and denominator coefficients of an IIR filter. Given that there is only a small number of coefficients (e.g. 8 each for numerator and denominator) there is little room for overfitting. This method is also effective to remove ringing associated with stimulus artifacts, for example when measuring responses to electrical stimulation (Oswal et al., 2016). In principle it is possible to average filter parameters estimates over events to get a more accurate estimate, however in practice the appropriate filter depends on the timing of the step relative to the sampling instants, which is unknown. The Steiglitz-McBride approach is easier for this reason.

In summary, a panoply of algorithms is available to address the issues raised in the Introduction. The next section illustrates their performance. These techniques target artifacts that are not easily removed with standard spatial filtering methods such as ICA, and thus are complementary with those methods.

## 3 Results

The algorithms are illustrated with synthetic data to clarify their properties, and with real data to evaluate their usefulness in a practical setting.

### Robust detrending

Figure 3 (bottom) shows the result of applying robust detrending rather than standard detrending. After a few iterations the algorithm has estimated the position of the glitch (grey line) and performed a weighted polynomial fit (red). subtracting the fit yields the detrended data (bottom right). Comparing to Fig. 3 middle right, it appears that the algorithm has achieved a better fit to the non-glitch parts, yielding a more useful detrended signal.

Figure 4 (top) shows one channel of real EEG data on which are superimposed 200 repetitions of a synthetic unipolar pulse with a shape similar to that shown in Fig. 2 (the low-amplitude pulses are not visible in the waveform at this scale). Averaging over trials reveals the pulse (black line in Fig. 4), but the linear trend that affects the raw data (Fig. 4, top) also affects the average over trials (Fig. 4, middle left). This can be addressed by applying a high-pass filter *before* cutting the data into trials, as is common practice in EEG data analysis. However doing so also seriously distorts the shape of the pulse (Fig. 4, middle right). Filtering *after* cutting into trials has an even worse effect due to onset response artifacts that repeat on each trial (not shown). As an alternative to high-pass filtering, the trial-averaged data may be detrended by fitting a low-order polynomial to the averaged data and subtracting the fit (Figure 4 bottom left). However, the presence of the pulse affects the fit (Fig. 4 middle left, red dotted line), adding a reverse trend to the response (bottom left). In contrast, the *weighted* fit (Fig. 4 middle left, red full line) leads to a better estimate of the target signal (bottom right).

**Figure 4:**
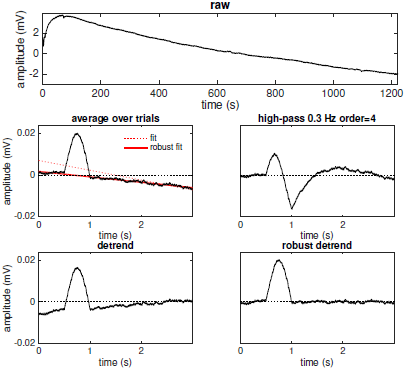
Robust detrending of EEG data. Data consist of 200 repetitions of a synthetic unipolar pulse of duration 500 ms and amplitude 20 μV superimposed on a real EEG signal (top). Middle left: trial average (black), linear fit (dotted red) and robust linear fit (full red) to trial average. Middle right: trial average of high-passed data (0.3 Hz cutoff, order 2). Bottom left: detrended trial average. Bottom right: robust detrended trial average. Data are from the same dataset as Fig. 1.

As an additional example, Fig. 5 (top left) shows one channel of raw EEG with multiple glitches, together with a robust polynomial fit of order 30 (red line). Figure 5 (top right) shows the detrended signal. The time scale of the variations that are included in the trend depends on the order of the polynomial, a higher order allowing finer fluctuations to be removed (see Discussion for caveats). As a final example, Fig. 5 (bottom left) shows a synthetic signal corrupted by both a 50 Hz sinusoidal artifact and a temporally-localized glitch (amplitude 100, shown truncated). An effective way to remove the sinusoidal artifact is to fit a quadrature pair of 50 Hz sinusoids (in lieu of “trend”) and subtract the fit, equivalent to removing that component in the Fourier domain, but the presence of the glitch disrupts this process and leads instead to a *greater* sinusoidal artifact amplitude (Fig. 5, bottom center). In contrast the weighted fit produces a clean signal (bottom right). Notch filtering is prone to similar effects, that weighted detrending conveniently avoids. This last example shows that robust detrending is not restricted to slow trends.

It is worth remarking that the same algorithm can be used instead to robustly follow the *slow* variations in the signal, analogous to a low-pass filter. The advantage over a standard lowpass filter is again that the result is not affected by localized glitches. These persist of course in the “smoothed” signal, but the shape of the non-glitch portions is not affected.

**Figure 5:**
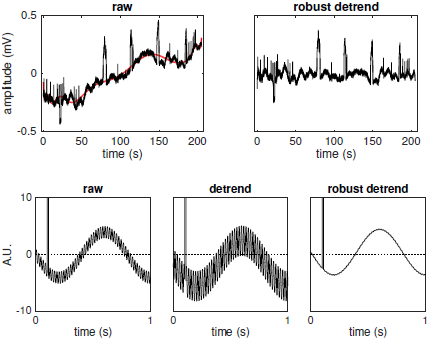
Top left: raw EEG signal (black) and 30th order polynomial fit (red). Top right: detrended signal. Data are from the same dataset as Figure 1. Bottom: Robust removal of a 50 Hz sinusoidal trend. Left: 1 Hz sinusoidal “signal” corrupted by 50 Hz artifact and a temporally-localized glitch (amplitude 100). Center: a 50 Hz sinusoidal function is fit to the data and subtracted. Right: same, but the fit is weighted by a weighting matrix that is zero at time of the glitch.

### Inpainting

Figure 6 (top) shows a 50-channel signal [*x_n_*(*t*)] produced by mixing 10 uncorrelated “sources”, consisting of sinusoids of different frequencies, via a 10 × 50 random mixing matrix. The signal of each channel was corrupted by adding a randomly-placed “ glitch” of duration 0.2 s. The positions of these glitches are indicated by the weighting matrix *[w_n_*(*t*)] (Fig. 6 middle). Applying the inpainting algorithm produces a cleaned signal (Fig. 6 bottom) indistinguishable from the glitch-free original (not shown). To achieve this result, the algorithm estimated the correlation structure of intact portions (*w_n_*(*t*) ≠ 0), and used them to design a set of projection matrices to reconstruct corrupt portions (*w_n_*(*t*) = 0). In this example there were 146 distinct configurations of intact/corrupt channels, and thus the same number of projection matrices. Note that none of the channels was glitch-free, and that many glitches overlapped in time.

**Figure 6:**
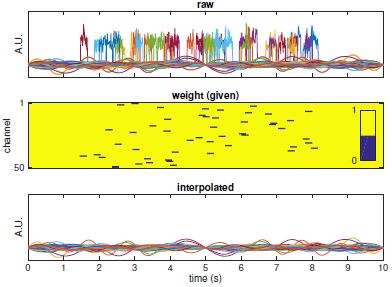
EEG signal inpainting. Top: 50-channel synthetic signal (rank 10) corrupted with randomly-placed “glitches” (thin lines). Middle: weighting matrix, zero at the positions of the glitches. Bottom: signal interpolated by the inpainting algorithm.

### Outlier detection

In the previous example, the weighting matrix *w_n_*(*t*) was known in advance. If no prior knowledge is available, a weighting matrix can be estimated from the data based on the assumption that glitches are uncorrelated across channels. Figure 7 (top) shows the weighting matrix estimated by the outlier detection algorithm (threshold = 1, 10 iterations) from the data plotted in Figure 6 (top). Based on this estimate, derived blindly from the data, the data can be effectively denoised (Figure 7 bottom), yielding a result almost identical to the original (not shown).

**Figure 7:**
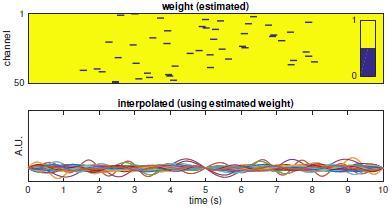
Outlier detection. Top: weight mask estimated from the signal plotted in Figure 6 (top). bottom: signal interpolated using the estimated weighting mask.

As another example, a segment of 128-channel EEG data was first detrended and then processed with the outlier detection algorithm (threshold =1,6 itera-tions). The estimated weight matrix is displayed in Fig. 8 (top). The pattern of outliers is quite diverse, with some channels affected most of the time (e.g. channel 39) and many channels some of the time (e.g. circa 2-3 s). Fig. 8 (middle) shows the time course of channel 35 before (black) and after robust detrending (green), and after outlier detection and inpainting (red). The signals are offset vertically for clarity. The invalid portion (between 2 and 4 s), visible as a horizontal line in the weights plot, is replaced by a combination of intact channels. Fig. 8 (bottom) shows similar data for channel 39 (weights visible as a horizontal dotted line across the full data segment). Here the algorithm has replaced a large number of samples, yielding an apparently cleaner signal (red trace).

**Figure 8:**
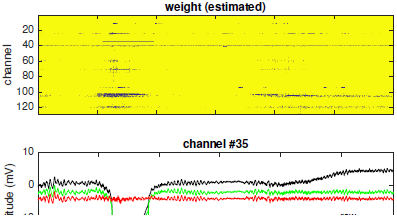
Outlier detection. Top: weight mask estimated from a segment of 128-channel EEG (detrended). Middle: one channel of EEG, showing the raw, de-trended, and interpolated signal, offset vertically for clarity. Bottom: same for another channel. Data are from the same dataset as Figure 1.

### Robust rereferencing

Figure 9 (blue) shows the time course of one channel of a segment of 128-channel EEG data. Another channel (not shown) was affected by an artificial glitch of amplitude -1000 mV, with the result that the channel average also shows a glitch (of amplitude ~-8mV) that is injected into the rereferenced waveform (Fig. 9, black). Robust rereferencing avoids this problem (red).

**Figure 9.**
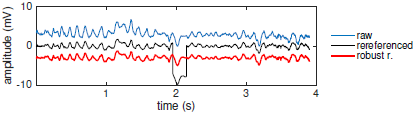
Robust rereferencing. Single EEG channel before (blue) and after standard rereferencing (black) and robust rereferencing (red).

### Step removal

Figure 10 (left) shows a sample of MEG data recorded with a phantom dipole source in an experiment that simulated conditions characteristic of deep brain stimulation (Oswal et al., 2016). Stimulation produces a high-amplitude magnetic pulse that can cause the electronics controlling the SQUID sensors to switch state, resulting in a sharp step of large amplitude. Such steps also occur spontaneously in MEG data (Gross et al., 2013). In this example, at each stimulus the MEG signal (black) is affected by a step that the algorithm removes (red).

### Ringing removal

Visible in the previous example are a series of glitches due to the ringing response from the low-pass antialiasing filter, that remains after step removal (Fig. 10, right). That ringing response (black) is effectively removed by the ringing removal algorithm (red). Ringing artifacts from other events (e.g. stimulus artifacts) can also be removed if their timing is known.

**Figure 10:**
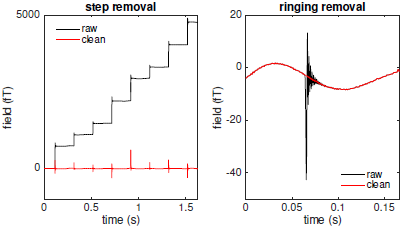
Step and ringing artifact removal. Left: one channel of MEG data in response to simulated deep brain stimulation (Oswal et al., 2016) (black) and the same signal after automatic step removal (red). Right: one channel of MEG from same data set showing a stimulus artifact (black) and the same signal after ringing removal (red). Data were recorded on a 275-channel CTF system at a 2400 Hz sampling rate (Oswal et al., 2016).

## 4 Discussion

### 4.1 The importance of preprocessing

Artifacts are deleterious in four ways. First, they *mask* interesting activity; for example, line noise may mask oscillatory cortical activity near 50 or 60 Hz, and electrode drift may mask slow cortical potentials. Second, they may *masquerade* as brain activity; for example a slow trend might lend a ramp-like shape to an evoked response, or ocular or myogenic artifacts might masquerade as cortical activity if they correlate with stimulation or behavior (Yeung et al., 2004; Yuval-Greenberg et al., 2008). Third, the need to attenuate them may prompt the use of processing such as high-pass filtering that entails deleterious *side effects.* Fourth, the artifacts may *impede analysis* by interfering with the processing; for example, high-amplitude artifacts may dominate the sums of squares involved in methods such as CSP, ICA, CCA, etc. Artifact removal is a prerequisite for good data analysis, and this paper offers a panoply of methods for the purpose. They are presented here together (rather than in separate papers) because they share concepts and processing, and because the interactions between methods need to be considered.

### 4.2 In which order?

Should a slow trend be removed before glitches, or after? Should they both be removed before spatial filtering, or after? At what stage should temporal filtering be applied, if at all? Linear operations may be swapped, but the data-dependent algorithms that determine their parameters depend on the order in which they are applied. Algorithms that rely on minimizing a sum of squares are sensitive to components with large variance: a trend or glitch may entice an algorithm to model it rather than sources of interest, and thus removing the artifact may make the algorithm more effective. Definitive guidelines are hard to set because of the variety of situations. As a rule of thumb, if algorithm *B* is sensitive to an artifact that algorithm *A* can remove, then *A* should be applied before *B*. A difficulty arises of course if *A* is also sensitive to artifacts that *B* can remove.

A typical EEG recording might be contaminated by a combination of *slow drift* specific to each electrode, *temporally localized glitches* also specific to each electrode, *eye blinks* shared across electrodes, *50 Hz and harmonics* shared across electrodes, *myogenic activity* that may be either electrode-specific or shared, *alpha activity* shared across electrodes, and so-on. A likely sequence might be: (a) discard pathological channels for which there is no useful signal, (b) apply robust detrending to each channel, (c) detect and interpolate temporally-local channel-specific glitches, (d) robust re-reference, (e) project out eye artifacts (e.g. using ICA or DSS), (f) fit and remove, or project out, 50 Hz and harmonics, (g) project out alpha activity, etc., (f) apply linear analysis techniques (ICA, CSP, etc.) to further isolate activity of interest.

A typical MEG recording might be contaminated by a combination of *SQUID jumps* specific to each sensor, *slow components* due to nearby vehicles and machinery (e.g. elevator), shared across sensors, *50 Hz and harmonics* from power lines and devices, shared across sensors, *sensor noise* specific to each sensor, *vibration artifacts* shared across sensors, *alpha activity* shared across sensors, *stimulus artifact*, etc. A likely sequence might be: (a) remove squid jumps, (b) remove stimulus artifacts, (c) remove associated antialiasing filter ringing artifacts, (d) isolate and remove slow environmental components, 50 Hz and harmonics, vibration artifacts, alpha activity, etc. using projection techniques or spatial filtering based on ICA, DSS, etc., (e) suppress sensor noise using the Sensor Noise suppression (SNS) algorithm (de Cheveigné and Simon, 2008), (f) apply linear analysis techniques (ICA, DSS, CSP, etc.) to further isolate activity of interest.

### 4.3 Comparison with other tools

A standard practice is to *discard* data corrupted by an artifact, for example discard a channel or trial. This is justified if those data are completely invalid, but wasteful otherwise. Data loss may be unacceptable if artifacts are spatially and/or temporally dense. One motivation for this work is to minimize such data loss while still attaining high data quality. Another is to avoid manual intervention, in contrast to common practices of *visual inspection* and *manual editing*. This goal is not fully attained, as it may still be necessary to manually adjust the order of processing stages or their parameters.

*Temporal filtering* is widely used (high-pass to remove drift, lowpass to attenuate sensor noise and myogenic artifacts, notch to remove power line components, etc.) but prone to the pitfalls noted in the Introduction. An aim of this work is to offer effective alternatives. *Detrending*, a popular alternative to high-pass filtering, is impaired by glitches. Robust detrending reduces this problem, giving the method a major advantage over filtering. Like non-causal filtering, detrending can raise causality concerns as the trend removed at time *t* reflects data that occurs after *t*. In practice the strong constraint imposed by the basis functions (e.g. smooth low-order polynomials) rules out the most egregious effects. Polynomial trend removal is implemented in toolboxes such as FieldTrip (Oostenveld et al., 2011), but without weighting. An alternative to fitting a polynomial is local linear regression, in which a linear fit is repeatedly applied to short segments that are moved forward in time, and the fits averaged. A robust version of this process is proposed in the `matlabmk` toolbox (http://kutaslab.ucsd.edu/matlabmk_fn_docs/matlabmk/robust_locdetrend.html). The robust rereferencing scheme of Bigdely-Shamlo et al. (2015) detects and interpolates corrupt channels over their entire duration, in contrast to the method reported here that does so only for corrupt segments of each channel.

A *spatial filter* is another common tool, with coefficients that are either predetermined (e.g. rereferencing, or Laplacian) or calculated from the data (e.g. ICA, PCA, etc.). A spatial filter allows multiple noise sources to be cancelled, but this works only if the dimensionality of the noise space (number of distinct active sources) is less than that of the data (number of channels). Usually, the coefficients of the filter remain constant over the data set, in other words the filter is *time invariant*. In contrast, the inpainting and outlier detection algorithms described here apply distinct spatial filters over different subsets of the time axis, similar to the STAR algorithm (de Cheveigné, 2016). The ability to apply different filters at different times sets these methods apart from standard time-invariant spatial filter methods such as PCA, ICA, CSP, beamforming, etc.

ICA is reported to benefit from high-pass filtering (Debener et al., 2010; Win-kler et al., 2015), and should thus benefit from detrending. More generally, the methods presented here can contribute to the success of component analysis in two ways. First, by taking care of high-variance components that would otherwise dominate the least-squares solutions involved in many methods, and second by taking care of numerous channel-specific artifacts that inflate the number of “sources” that need to be resolved from the limited number of sensors available (Debener et al., 2010). It should be noted that the methods presented here do not usually reduce the rank of the data. They thus complement standard methods such as ICA.

The combination of outlier detection and inpainting can be seen as a “robust” version of our earlier SNS algorithm which projects each channel on the subspace spanned by the others (de Cheveigné and Simon, 2008). It can also be seen as a generalization of our earlier STAR algorithm which only allows one corrupt channel at any time (de Cheveigné, 2016). It is similar in spirit to the Artifact subspace Reconstruction (ASR) method (Kothe, 2013), and related to spatial interpolation methods such as Perrin et al. (1989) that require neighboring channels to be intact. Only between-channel correlation is exploited here, for inpainting and outlier detection; an alternative is to exploit *temporal* structure within each channel via an autoregressive or wavelet model, or both space and time via a multichannel autoregressive model (Lawhern et al., 2012). Many methods have been developed for image, audio and video inpainting and matrix completion (Candès and Recht, 2008) that may also be of use for electrophysiological data. These remain to be fully explored.

The standard approach to address SQUID jumps in MEG is to discard corrupted data (Gross et al., 2013), and popular toolboxes such as FieldTrip (Oostenveld et al., 2011) or Brainstorm (Tadel et al., 2011) offer tools to detect such jumps. Discarding is not an option if the steps are dense on a majority of channels (e.g. Oswal et al. (2016)). In contrast, the method described here removes the steps with no data loss. After steps are removed, *ringing artifacts* due to the antialiasing filter remain, and the same artifacts may be observed in both EEG and MEG in response to electric stimulation (cochlear implant, deep brain stimulation, TMS, etc.). The standard approach is to discard the ringing interval together with the artifact (Herring et al., 2015), but this is not an option if artifacts are dense. As far as we know, our methodology has not been reported elsewhere.

Robust techniques are well developed in statistics and data mining (Aggarwal, 2016), and some have been applied to EEG or other signals (Pernet et al., 2011; Lepage et al., 2014; Bigdely-Shamlo et al., 2015; Krauledat et al., 2007). Common to these techniques is the weighting of the data according to their reliability. The weighting is often implicit in the definition of the robust statistic, or in the estimation process, in contrast to the explicit weighting matrix that we use here. An advantage of explicit weights is that a measure of reliability derived in one context (e.g. detrending) can be applied in another (e.g. rereferencing).

*Outliers* are detected here based on mismatch from a model (spatial or temporal), rather than on the basis of their extreme values as is common. To derive the model we use standard statistics such as the mean and standard deviation. More robust statistics (median, etc.) have been advocated (Leys et al., 2013), but using them is unlikely to make a large difference in this context. Binary weights are used throughout; the use of graded weights is supported by the implementation, but unlikely to offer a major benefit.

### 4.4 Caveats and failure scenarios

#### Detrending

Assumes a signal model (basis) that must be flexible to fit the trend, yet inflexible so as to not absorb fluctuations of interest. The choice of parameters (e.g. polynomial order) is critical to correctly set the time scale of the trend to be removed, and success in this respect is uncertain if target and trend share a similar time scale. *Robust* detrending can fail if a glitch is mistaken as a trend (for example if it has a long temporal extent), or a trend mistaken as a glitch. To avoid this, it may be useful to first detrend using a low-order model, and then a higher-order model with weights constrained by the first fit. Simple trends are easily fit with a low order polynomial, or the first few terms of a Fourier series. Fitting more complex trends may run into difficulties due to what is known as Runge’s effect (polynomials) or Gibb’s effect (sinusoids) (Platte et al., 2011). As an example, in Figure 5 (top right) the 30-th order polynomial fit leaves a residual fluctuation that is not completely eliminated by a higher-order order fit (not shown). A potential solution is to use splines (Shumway and Stoffer, 2011), or weighted smoothing interpolation (Davies and Meise, 2008), but these solutions have not been explored here. A simpler expedient is to apply detrending to shorter segments for which a low-order model is adequate. The outcome of *robust* detrending is dependent on the choice of the threshold parameter (in addition to those of the trend model), and we know of no convergence proof.

#### Inpainting

Fails if the signal to reconstruct is not contained in the subspace spanned by the intact channels (signal model 2). This can happen if the number of sources active at that time is greater than that the number of intact channels. In the extreme, if there are *no* intact channels, the algorithm cannot proceed (in that case the data is left untouched). Inpainting may also fail if there are insufficient data to reliably estimate the reconstruction matrices. Inpainting also fails if the channel is entirely corrupt, as the projection matrices cannot be estimated. The best option is to discard the channel (and possibly interpolate from intact neighbors).

#### Outlier detection

Assumes that the intact data match model 2, outliers being defined as samples that do not match the model. A same data set might match multiple models. For example two partly overlapping glitches could be interpreted as a pulse common to both (the part that for which they overlap), together with shorter glitches specific to each. The algorithm can chose either, and the choice that it makes might be unexpected. There is no guarantee of convergence of the outlier detection, and indeed the solution is often observed to fluctuate. That said, the algorithm typically gives plausible and useful results on real data.

#### Robust rereferencing

Is contingent on the quality of the outlier detection that produces the mask, although the impact of an error on any particular channel is mitigated via the averaging process.

#### Step removal

Could in theory remove genuine step-like activity, but in practice there is never any ambiguity and the method is robust. It runs into difficulty if the sections between steps are not flat. In that case a combination of step removal and detrending may be required.

#### Ringing removal

Treats a short segment of data as a filter impulse response, that it then removes. Brain signal features that occur during that short segment might be removed as well, although the likelihood of this happening is limited by the constraints imposed by the low-order IIR filter. The risk could be further reduced by fitting all ringing artifacts with exactly the same filter, although this option was not explored.

The methods described in this paper deal only with *channel specific* artifacts; artifacts that impinge on multiple channels are not addressed at all.

### 4.5 Implementation

The routines are implemented in Matlab within the NoiseTools toolbox (http://audition.ens.fr/adc/NoiseTools/). They are based on closed-form solutions, possibly iterated a small number of times, and thus most methods are cheap in terms of computation. Inpainting and outlier detection are somewhat more costly because a large number of combinations of valid/invalid channels may need to be considered (potentially up to 2*^N^* for *N* channels, in typical practice a few thousand). It can happen that channels within 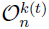 are rank-deficient over 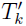; to handle this situation, for each k those channels are submitted to a PCA, the PC series is truncated, and the projection is performed on the remaining PCs. The eigendecomposition required by this step is the main computational bottleneck. To limit the cost of eigendecomposition (*O*(*N*^3^)), for each channel *n*, a subset of size *M* < *N* of channels “closest” to channel *n* is selected on the basis of physical proximity or correlation with *x_n_*(*t*). To further reduce computation, configurations that involve only a few data samples are skipped, and the corresponding data are reconstructed by serial interpolation as a weighted sum of neighboring intact samples.

The algorithms are described here in terms of batch processing, but the statistics that they require (mean, covariance, cross-correlation) can be calculated incrementally and updated in real time, potentially allowing the methods to be deployed in a real-time brain-computer interface (BCI) system. One such application, cognitive control of a hearing aid, is a primary motivation of this work.

## 5 Conclusion

This paper presents a set of methods to preprocess multichannel data such as EEG or MEG and improve their quality. These methods address ubiquitous sources of artifact that corrupt data and interfere with analysis and interpretation, and complement other methods such as temporal or spatial filtering, either as a replacement with better performance and fewer drawbacks, or as a complementary processing step to make them perform better. The methods include robust detrending, robust rereferencing, outlier detection, inpainting (interpolation), step detection and correction, and ringing artifact removal.

## Acknowledgements

This work was supported by the EU H2020-ICT grant 644732 (COCOHA), and grants ANR-10-LABX-0087 IEC and ANR-10-IDEX-0001-02 PSL^*^. Vladimir Litvak kindly pointed us to the MEG data of the study of Oswal et al. (2016) made available at https://figshare.com/articles/phantom090715_BrainampDBS.20150709_01_ds_zip/4042911/3, and helped test the step removal and ringing removal algorithms.

